# Transmission of *Xylella fastidiosa* subsp. pauca ST53 by the sharpshooter *Cicadella viridis* from different source plants and artificial diets

**DOI:** 10.1101/2022.10.25.513644

**Authors:** Nicola Bodino, Vincenzo Cavalieri, Maria Saponari, Crescenza Dongiovanni, Giuseppe Altamura, Domenico Bosco

## Abstract

The sharpshooter *Cicadella viridis* L. (Hemiptera: Cicadellidae) is the most common sharpshooter in Europe and, given its xylem feeding behaviour, is considered a potential vector of the plant pathogenic bacterium *Xylella fastidiosa* Wells et al. (Xanthomonadales: *Xanthomonadaceae*). We tested *X. fastidiosa* subsp. pauca ST53 (*Xfp*) transmission capabilities of *C. viridis* adults, namely i) acquisition efficiency from four host plant species – periwinkle, milkwort, lavender, alfalfa – and from two artificial diets (PD3 and Xfm), ii) inoculation efficiency to periwinkle at different times post acquisition from different plant and artificial diet sources. The main European vector species – *Philaenus spumarius* – was used as a control. *Cicadella viridis* was able to acquire *Xfp* from periwinkle, milkwort, and lavender, although with low efficiency (3–16%) and from artificial diets (23–25%). Successful inoculation on periwinkle was extremely rare, being observed only three times, following feeding on milkwort plant and PD3 artificial diet sources. Our study shows that *C. viridis* is not a relevant vector of *Xfp*, given the very low transmission rate in controlled conditions and the inability to feed on olive. The low efficiency reported here sums also to ecological constraints (mainly monocots host plants, humid environments) that make difficult to forecast a relevant role in dispersing *X. fastidiosa*, at least within the present distribution of the exotic bacterium in Europe. However, a possible role of this species in spreading *Xf* in other agroecosystems, e.g. vineyard and stone fruits grown in humid areas, cannot be excluded.

## Introduction

The discovery of the xylem-limited bacterium *Xylella fastidiosa* (*Xf*) in Europe, and its associated epidemics on different crops (“Olive Quick Decline Syndrome” (OQDS) on olive in Apulia (southern Italy), “Almond Leaf Scorch” (ALS) on almond in Spain) (Saponari et al. 2017, Moralejo et al. 2019), have boosted research efforts on insect vectors throughout the Continent (EFSA Panel on Plant Health (PLH) 2015, Cornara et al. 2018). The key vector species in the Mediterranean region had been soon identified within the xylem-sap feeders spittlebugs (Hemiptera: Aphrophoridae): *Philaenus spumarius* L. and, to a lesser extent, *Neophilaenus campestris* (Fallén) and *Philaenus italosignus* Drosopoulos & Reman (Cornara et al. 2017, Cavalieri et al. 2019). However, species belonging to another group of xylem-sap feeder – the sharpshooters (Hemiptera: Cicadellidae: Cicadellinae) – are considered as potential vectors in Europe since they represent the predominant *Xf* vectors in the Americas (Redak et al. 2004, Almeida et al. 2005, Cornara et al. 2019, EFSA Panel on Plant Health (PLH) et al. 2019). Compared to the New World, sharpshooters are underrepresented in the Palearctic region, but at least one species, *Cicadella viridis* (L.), is quite widespread and locally abundant in Europe (Nickel 2003, Pavan and Gambon 2004, De Jong et al. 2014, EFSA Panel on Plant Health (PLH) et al. 2019). This species is unlikely to be present in the areas currently facing *Xf* outbreaks in the Mediterranean region, being generally associated to humid meadows and grasslands, but it is considered a relevant vector candidate in North Italy and Central Europe, especially on grapevine (Janse and Obradovic 2010, EFSA Panel on Plant Health (PLH) et al. 2019, Hasbroucq et al. 2020, Markheiser et al. 2020). However, this species can be present in some olive agroecosystems, and was repeatedly collected in the ground cover of olive orchards in both Northern Italy — Veneto Region (Nicola Mori and Domenico Bosco, personal observations) — and Southern Italy — Abruzzo (Sanna et al. 2019) and Basilicata Region (Trotta et al. 2021).

To the best of our knowledge, no information on the capability and efficiency of *C. viridis* to transmit European *Xf* strains are available yet. Given the relevance of the OQDS epidemics, in this study we investigated the competence of *C. viridis* to transmit *Xf* isolates of the subsp. pauca (*Xfp*) and harbouring the sequence type 53, found to be the causal agent of the OQDS. To estimate its potential role as a vector, we performed different microcosms assays to study i) the acquisition efficiency and persistence of *Xf by C. viridis* from different sources (four plant species and two artificial diets), and ii) inoculation efficiency on recipient plants at different times post acquisition.

## Materials and Methods

Transmission experiments were carried out for three consecutive years, from 2017 to 2019, in the demarcated infected area of the Apulia region (Southern Italy). Transmission tests were performed using the olive infecting strain De Donno (subsp. pauca, sequence type ST53).

### Insects and plants

*Cicadella viridis* nymphs and adults were collected in wet meadows located near Torino in the Piedmont Region of Italy, one to two weeks prior to the start of each experimental assay. Insects were maintained in mesh and plastic fabric cages (Bugdorm: 75×75×115 cm) containing potted host plants (*Avena sativa* L.*, Sorghum halepense* L. (Pers)). The cages were placed in a climatic chamber (24.7 ± 1.0 °C, 72.1 ± 7.2% RH) in the “Li Foggi” nursery of ARIF (Agenzia Regionale Attività Irrigue e Forestali) (Gallipoli, province of Lecce). The air temperature and relative humidity were recorded hourly under both conditions using data loggers (HOBO U23-002; Onset Computer, Bourne, MA, USA). To confirm the *Xfp-free* status of the reared sharpshooters, 20–30 individuals were randomly collected from maintenance cages before starting each acquisition assay and individually tested by quantitative real time PCR (qPCR) (Harper et al. 2010). In all transmission tests, adults of *Philaenus spumarius* were included as control. These latter adults were collected in olive groves located in *Xf*-free areas in northern part of the Bari province, and maintained on caged plants of *Sonchus oleraceus* L. and *Vicia faba* L., under the same conditions of *C. viridis*. Quantitative real time PCR (qPCR) assays were performed on randomly sampled individuals to confirm they were negative to *Xfp*.

Preliminary tests performed in summer 2017 showed high mortality of *C. viridis* adults when caged on *Xfp*-infected olive plants, indicating that this species is unable to feed on olive, and therefore this host plant was not used as source plant in the subsequent transmission tests.

Source plants used for the acquisition included: periwinkle (*Catharanthus roseus* (L.) G.Don) in autumn 2017, lavender (*Lavandula angustifolia* Mill.) in summer 2018, myrtle-leaf milkwort (*Polygala myrtifolia* L.) in autumn 2018, alfalfa (*Medicago sativa* L.) in summer 2019. All these species were proven to be both susceptible to *Xfp* ST53 and suitable hosts for *C. viridis* and *P. spumarius*. To obtain systemically infected source plants, needle-inoculations were performed several months in advance, using bacterial suspensions prepared by scraping 7-10 days old *Xfp* ST53 colonies grown on PD3 medium. More specifically, young plantlets of periwinkle and alfalfa were inoculated 3 months before performing the experiment; lavender and myrtle-leaf milkwort plants were inoculated on multiple shoots at least 6-8 months before the transmission experiments. Inoculated plants were maintained in confined quarantine greenhouse at controlled temperature of ranging between 22°C and 28°C. After the inoculation, leaf tissues were sampled and tested by qPCR (Harper et al., 2010) to assess bacterial multiplication and host colonization. Only plants showing qPCR-positive reactions for the leaves sampled distantly, 10-15cm far, from the inoculation points were considered systemically infected and retained for the acquisition experiments. Indirect estimation of the bacterial load in these source plants, by interpolation of the quantification cycles (Cq) on a standard calibration curve, indicated a population size in the range of 10^6 CFU/ml (i.e. corresponding to Cq of 22-24).

### *Xf* transmission from source plants

Four independent transmission tests were performed to assess the acquisition and transmission efficiency of *Xfp* ST53 by adults of *C. viridis*, as well as the bacterium retention and multiplication within the sharpshooter foregut. Insects were collected from maintenance cages in batches of 150-300 individuals and isolated on three source plants within a mesh/plastic cage (Bugdorm: 75×75×115 cm). Acquisition period (AAP) lasted for two days (48h). Adults of *P. spumarius* were used as control in all the assays, hence batches of 150-300 spittlebugs were caged onto the same source plants for the same AAP as *C. viridis* individuals.

At the end of each AAP, individuals from both insect species were transferred to new cages containing potted plants of non-host species of *Xfp* ST53, (same as described above for maintenance). Insects were randomly collected from the maintenance cages at different times after the end of AAP (0 – 3 – 7 – 14 days) and transferred onto recipient plants (periwinkle, three-month old) for an IAP of three days (72 h). Five insects were isolated on each periwinkle, enclosed in a mesh sleeve, with access to the entire plant. Five replicas per each insect species and each timepoint after the end of AAP were carried out. Because of the high mortality rate registered during and after AAP on some plant species, *C. viridis* individuals were not always tested at the further time points and/or a lower number of replicas was performed at each time point (see results below). The duration of both AAP and IAP were chosen from preliminary trials and previous studies to maximize survival of the sharpshooter. Both IAP and post-AAP rearing took place in climatic chamber in the previously mentioned location (Vivai “Li Foggi” ARIF) at the same conditions described above.

At the end of each IAP, alive insects were collected by mouth aspirator, and individually stored in ethanol 90% at −20 °C. Insects survival rate was recorded for each AAP and IAP assay. Recipient plants (periwinkles) were treated with a systemic insecticide (Imidacloprid) and kept in a climatic chamber at 25 ± 2 °C. Inoculated periwinkles were analysed by qPCR (Harper et al., 2010) for the presence of *Xf* three months after the experimental assay.

### *Xf* transmission from artificial diet

In autumn 2018, *C. viridis* acquisition capability was tested in an artificial nutrition assay using the reference *Xfp* ST53 strain De Donno. The bacterial strain was grown on *Xf*m (minimum defined medium for *Xf*) and PD3 (Almeida et al. 2004) media. On *Xfm* medium, slow and limited bacterial growth was recorded, with colonies observed only after more than 3 weeks, even so, they were recovered and used for the experiment. Whereas, on PD3 agar plates colonies were observed and recovered after 8-10 days. Bacterial colonies were scraped from both media and suspended in a diet solution (L-glutamine, 0.7 mM; L-asparagine, 0.1 mM; sodium citrate, 1 mM; pH 6.4), with a final concentration adjusted to 4×10^8^ CFU/mL (OD_600_ 0.5).

The artificial diet containing the bacterial strain De Donno was offered to sharpshooters through a membrane feeding system. Each replicate consisted in a 4-cm diameter plastic cup, with 500-μL cell suspension loaded between two layers of stretched Parafilm located at the base, and with mesh at the opposite end. Insects were inserted in the cup from a hole (diameter 1 cm), which was then sealed with a cotton plug. Replicates were placed in random position on trays over a sheet of green cardboard to stimulate probing by the sharpshooters. We replicated the test 40 (PD3 thesis) and 8 (*Xfm* thesis) times, with five insects inserted in each cup, for a total of 200 (PD3) and 40 (*Xfm) C. viridis* adults tested. The AAP lasted 12h at 22 ± 2 °C. The inoculation assays followed the same procedure described above for the plant transmission assays. Insects were hence transferred in 5-individual batches onto periwinkles for a three-day IAP at different times after the AAP end (up to 14 days from AAP end). Insects were collected and analysed as described above.

#### Detection of *X. fastidiosa* in the insects

All alive individuals of *C. viridis* and *P. spumarius* collected from the cages after the IAP were singly tested by qPCR upon purifying the DNA from the excised heads using a CTAB-based extraction protocol (Cavalieri et al., 2019). The assays included three replicates of a 10-fold serial dilution of artificially spiked insect extracts with known bacterial concentrations ranging from 10^6^ to 10^1^ CFU/mL. An estimation of the bacterial concentration (CFU/head) in the positive specimens was extrapolated from the standard curve generated by quantitation cycle values of the spiked samples against the logarithm of their concentration.

### Statistical analysis

The proportion of *Xf*-positive insects and the proportion of infected recipient plants were modelled separately by bayesian logistic GLM (binomial link), with *Insect species, Source plant, Time post-acquisition*, and the interaction *Insect species* * *Source plant* as the fixed covariates. Bayesian GLM was used, instead of frequentist GLM, to appropriately deal with zero infection rates in alfalfa trial. *Xf* load in infected insects, i.e. the number of bacterial cells in insects’ mouthparts, was modelled by linear regression, with *Insect species, Source plant, Time post-acquisition*, and the interaction *Insect species* * *Source plant* as the fixed covariates. The size of the *Xf* population was decimal log transformed (log10(x + 1)) in all the tests to meet the model assumptions. No statistical models were performed on proportion of infected plants given the low number of *Xf*-positive plants after inoculation by *C. viridis*. Bayesian GLM were performed with *bayesglm* function in arm package. All analysis were performed in R 4.0.3 (R Core Team 2020), with arm (Gelman et al. 2021) and ggplot (Wickham 2016) packages.

## Results

### Survival of tested insects

The survival rate of *C. viridis* during AAP and post-AAP period was low for those fed on lavender and myrtle-leaf milkwort [≈ 20% (70/300) and ≈ 50% (150/300) respectively)], whereas it was generally higher on periwinkle (80–70%). No survival rate during AAP on alfalfa was registered due to logistic constraints. During the IAP, the survival rate of *C. viridis* was 78.8% (126 alive on 160 tested) and 70% (70/100) for those individuals previously isolated for AAP on periwinkles and myrtle-leaf milkwort, respectively. Survival rate was instead significantly lower for those previously isolated on lavender (32%, 16/50), and alfalfa (30%, 9/30). Also, all the sharpshooters fed on lavender died within 5 days from start of AAP. The control assays carried out with *P. spumarius* showed a high survival rate of individuals during IAP on all the tested plants, ranging from 100% on alfalfa to 67.4% on myrtle-leaf milkwort, with a significant difference between these two source plants only (Tab. 1).

**Table 1:** Parameters estimated from logistic GLM or linear regression of fixed covariates *Source plant, Insect species, Time post-acquisition* on the survival of insects during IAP, the proportion of infected individuals, and *Xylella fastidiosa* population size at the end of IAP.

|  | Survival rate during IAP |  | PCR-positive insects |  | *Xf* population (positive samples only) |  |
| --- | --- | --- | --- | --- | --- | --- |
| **(Intercept)** |  |  |  |  |  |  |
| *Cicadella viridis* on *Lavender* | -0.62<br>* | (-1.07, -0.19)<br>[-2.20] | -2.09<br>*** | (-2.10, -2.08)<br>[-3.88] | 5.46<br>*** | (3.61, 7.31)<br>[5.90] |
| **Source plant** |  |  |  |  |  |  |
| *Lavender* |  | — |  | — |  | — |
| *Alfalfa* | -0.17 | (-0.88, 0.56)<br>[-0.36] | -1.66 | (-1.67, -1.65)<br>[-1.58] |  | — |
| *Periwinkle* | 1.90<br>*** | (1.25, 2.48)<br>[5.64] | -0.75 | (-0.77, -0.74)<br>[-1.09] | -3.26<br>** | (-5.36, -1.15)<br>[-3.09] |
| *Milkwort* | 1.43<br>*** | (0.85, 2.00)<br>[4.09] | 1.17<br>* | (1.16, 1.18) [2.02] | -2.12<br>* | (-4.10, -0.13)<br>[-2.13] |
| **Insect species** |  |  |  |  |  |  |
| *Cicadella viridis* |  | — |  | — |  | — |
| *Philaenus_spumarius* | 2.22<br>*** | (1.77, 2.91)<br>[-5.89] | -0.52 | (-0.53, -0.51)<br>[-0.81] | 1.56 * | (0.32, 2.81)<br>[2.51] |
| **Insect species \* Source plant** |  |  |  |  |  |  |
| *Philaenus_spumarius* * *Alfalfa* | 3.07 * | (0.26, 5.89)<br>[1.99] | 1.78 | (1.77, 1.80) [1.50] |  | — |
| *Philaenus_spumarius* * *Periwinkle* | -2.55<br>*** | (-3.44, -1.81)<br>[-5.19] | 2.59<br>** | (2.57, 2.60) [3.09] |  | — |
| *Philaenus_spumarius* * *Milkwort* | -2.28<br>*** | (-3.01, -1.52) [-4.78] | 0.79 | (0.78, 0.80) [1.10] |  | — |
| **Time post-AAP** | — |  | -0.09<br>** | (-0.09, -0.09)<br>[-2.76] |  | — |
| N | 59 |  | 613 |  | 68 |  |
| logLik | -51.67 |  | -165.90 |  | -157.65 |  |

\*\*\* p < 0.001; \*\* p < 0.01; \* p < 0.05. Z statistics in brackets.

### Acquisition and transmission of *X. fastidiosa* by source plants

Acquisition efficiency varied significantly between insect species and source plants (*Insects species* × *Source plants*: 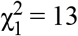, P = 0.004). *Cicadella viridis* was able to acquire *Xf* from three out of four source plants tested (periwinkle, lavender, myrtle-leaf milkwort). No *Xf*-infected insects were recollected from alfalfa source plants. The acquisition rate was low, although markedly different among the tested plant species (Fig. 1, Tab 1). The overall percentage of *Xf*-infected sharpshooters was significantly higher in those fed on milkwort (16.4%) compared to periwinkle and alfalfa (2.7–0%) (Tab 1). Acquisition rate from myrtle-leaf milkwort was quite high immediately after the end of AAP (50%, 5/10) and *Xf* was retained up to 17 days, even though at notably lower prevalence (≈ 10-15%). Conversely, only few individuals were *Xf*-positive after AAP on periwinkle (2.7%, 3/112) and Lavender (12.5%, 4/32). All the *Xf*-positive insects from these source plants were collected within 8 days from the start of AAP (Fig 1).

**Fig. 1:**
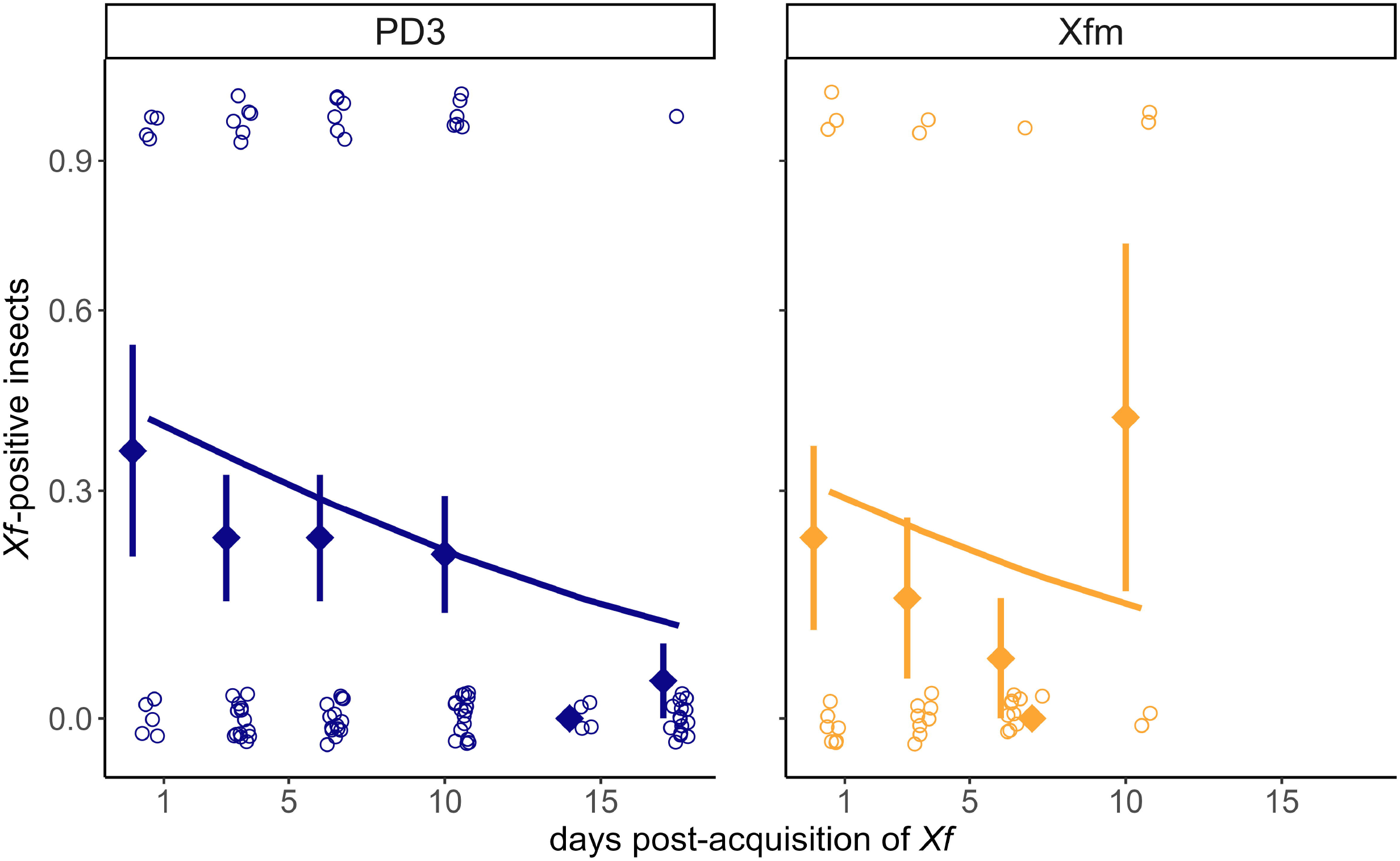
Proportion of *Xf*-positive *Cicadella viridis* and *Philaenus spumarius* individuals at different time points after acquisition from different source plant species. Hollow points represent the results of single insects (positive = 1, negative = 0); a small amount of variation is included to make visible the number of insects tested at each time point even if the multiple points had the exact same value. Red filled points and line represent the proportion of positive insects (mean ± SEM) and the black lines the outcome of bayesian binomial GLM model showed in Tab. 1.

*Philaenus spumarius* was able to acquire *Xf* from all the tested source plants, although with significant differences in efficiency (Tab 1). The proportion of *Xf*-infected spittlebugs was significantly higher in those fed on periwinkle (19.6%), and myrtle-leaf milkwort (21.6%) than the other two source plant species (lavender: 2.3%; alfalfa: 5.7%). The acquisition efficiency of *P. spumarius* was significantly higher than *C. viridis* only when the source plant was periwinkle (Tab 1).

Estimated *Xf* load was in average lower in *C. viridis* than *P. spumarius* infected individuals (median range *C. viridis:* 60-2000 cells, *P. spumarius:* 170-8300), and slightly lower in individuals from periwinkle [38 CFU/head (CI: 9–154)] compared to those from lavender source plants [683 CFU/head (CI: 109–4240)], regardless the insect species (*Insect species:* F_1,48_ = 5.22, P = 0.027; *Source plant:* F_2,48_ = 3.17, P = 0.05) (Supp. Figure S1). After acquisition from periwinkle source plants, the *Xf* load in the few *Xf*-positive *C. viridis* individuals rapidly declined to few *Xf* cells within one week post-AAP.

A single periwinkle recipient plant resulted positive to *Xf* after inoculation by *C. viridis* previously fed on milkwort, while no successful inoculations by insects fed on periwinkle, lavender, or alfalfa were obtained (Fig. 2). The infected plant was inoculated immediately after the end of AAP by a batch of sharpshooters encompassing two positive individuals. Conversely, *P. spumarius* was able to successfully inoculate several periwinkle recipient plants after AAP on *Xf*-infected periwinkles (46.7%, 7/15) myrtle-leaf milkwort (45.5%, 10/22), and *lavender* (9.1%, 2/22), but not after feeding on infected alfalfa.

**Fig. 2:**
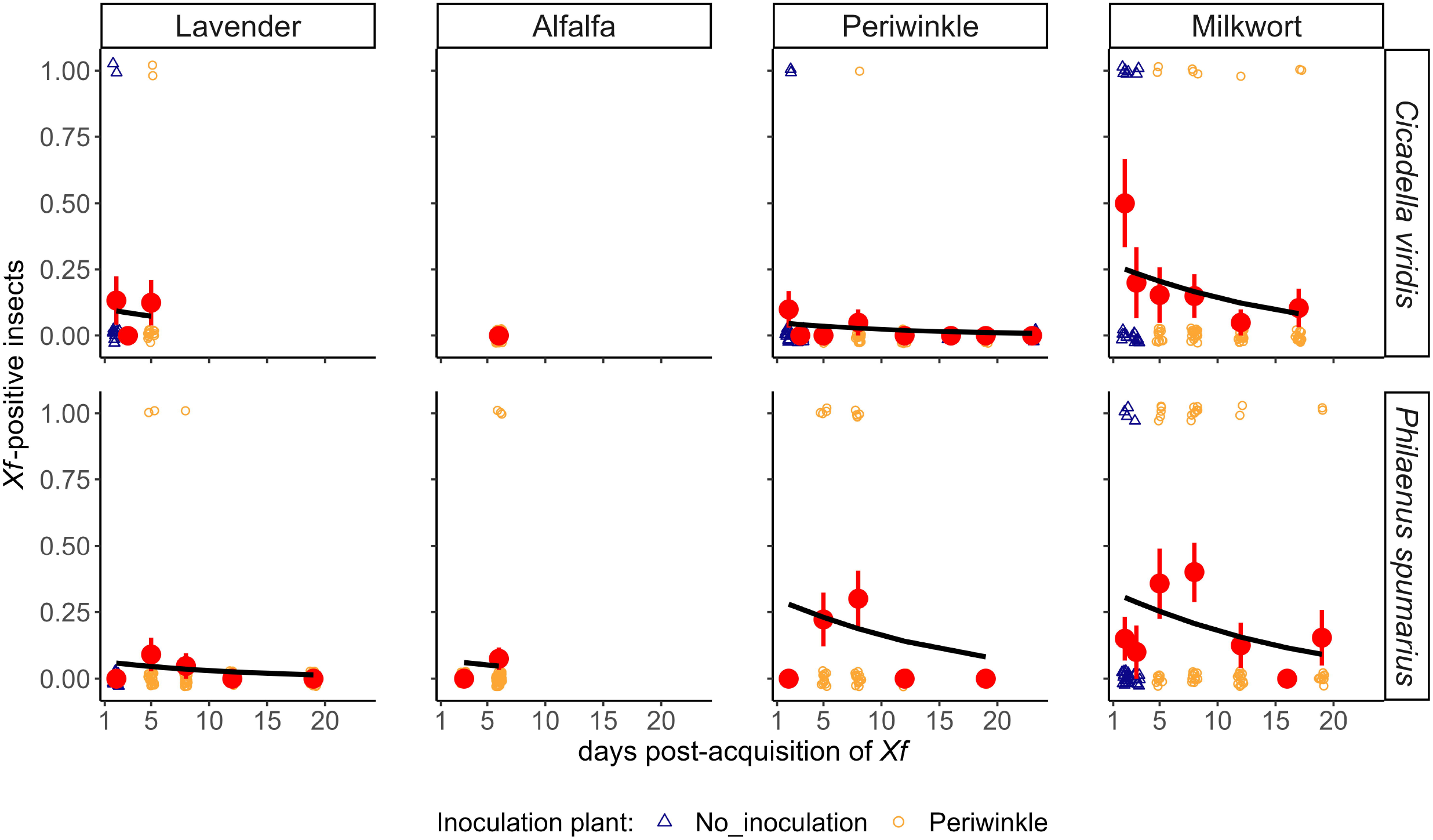
Proportion of *Xf*-positive periwinkles inoculated by batches (n = 5) of insects (*Cicadella viridis* and *Philaenus spumarius*) at different time points after acquisition from different source plant species. Hollow triangles represent the results of single plants (positive = 1, negative = 0); a small amount of variation was included to make visible the number of plants tested at each time point, even if the multiple points had the exact same value. Green filled points represent the proportion of positive olive plantlets (mean ± SE)

### *Xf* transmission from artificial diet

The mortality of *C. viridis* during AAP in the feeding arena was low (PD3: 5.5%, 11/200;*XF*M: 0%, 0/40). The overall *Xf* acquisition rate was not significantly different when the bacterium used to prepare the artificial diet was grown on PD3 or Xfm (PD3: 25.3%, 23/91; Xfm: 22.9%, 8/35), but the proportion of infected insects decreased steeply at longer post-AAP times for both diets (*Diet:* 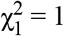, P = 0.309; *Time post-AAP*: 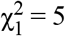, P = 0.025) (Fig 3). *Xf* load in sharpshooter foregut was low – without significant difference between the two diets – with estimated values mainly comprised between 10-100 cells, and only two individuals from PD3 diet with 200-300 estimated bacterial cells (*Diet:* F_1,28_ = 3.36, P = 0.08). Two out of 21 tested recipient plants (9.5%) were *Xf*-positive after inoculation by *C. viridis* fed on *Xf*-PD3. Both infected plants were from those inoculated 6-day post-AAP. No PCR-positive recipient plants inoculated with sharpshooters fed on *Xf-Xfm* were found.

**Fig. 3:**
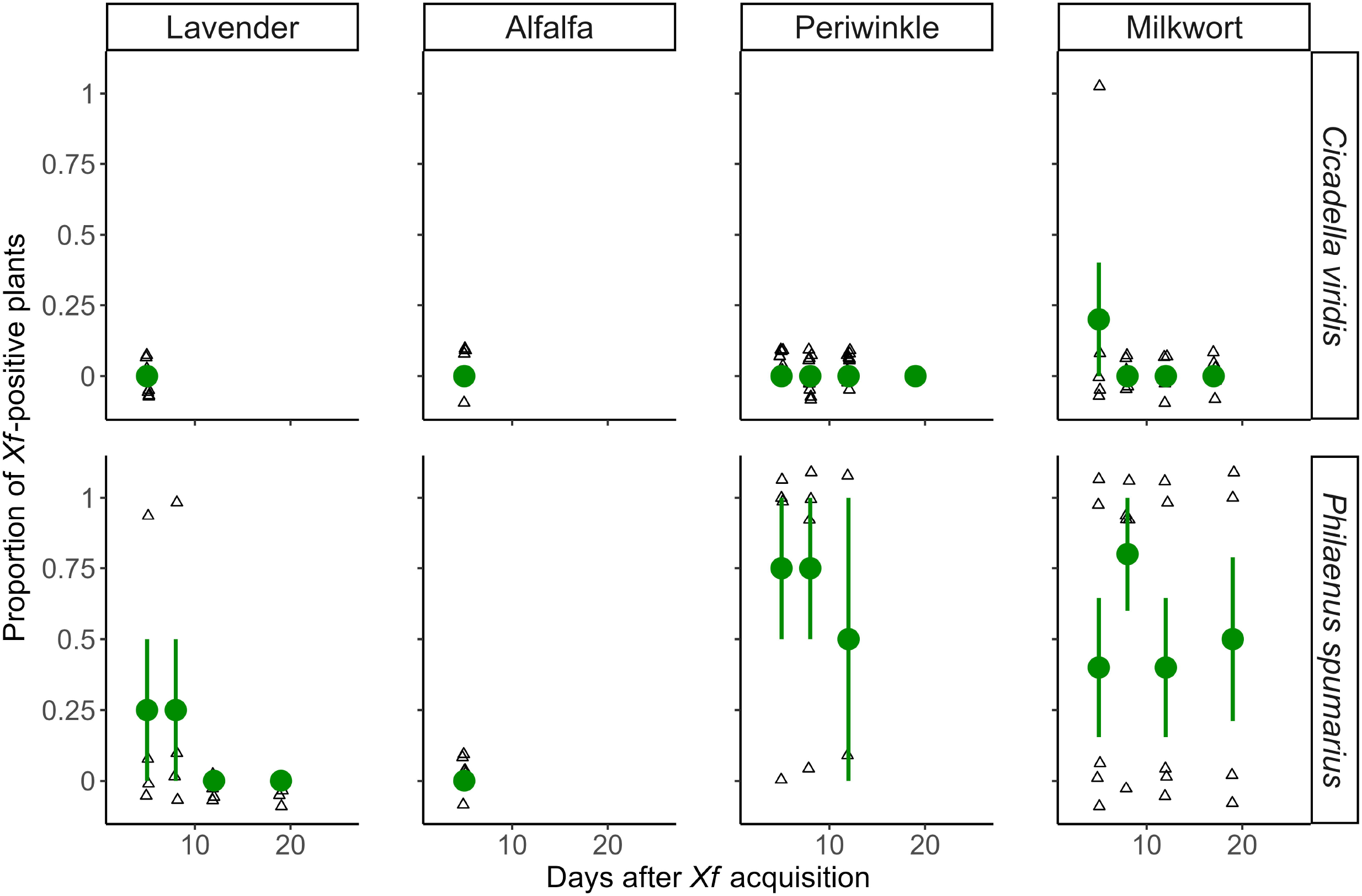
Proportion of *Xf*-positive *Cicadella viridis* individuals at different time points after acquisition from different artificial diets. Hollow points represent the results of single insects (positive = 1, negative = 0); a small amount of variation is included to make visible the number of insects tested at each time point even if the multiple points had the exact same value. Red filled points represent the proportion of positive insects (mean ± SEM) and the lines the outcome of binomial GLM model.

## Discussion

This study investigated the ability of the sharpshooter *C. viridis* in transmitting *Xf* ST53, in comparison to that of the main *Xf* vector in Europe, the spittlebug *P. spumarius*. Four source plant species and two artificial diets were tested in acquisition-inoculation assays. Olive as source plant was not included since preliminary trials showed that *C. viridis* do not survive on it.

*Cicadella viridis* was able to acquire *Xf* from three out of four source plant tested, and from both artificial diets. However, the acquisition efficiency was generally low (0-16%), as well as *Xf* population in its foregut. In comparison, acquisition rates of *P. spumarius* on the same source plants were higher on periwinkle and similar on the other source plants than those observed for *C. viridis*. Inoculation success rate was considerably lower for *C. viridis* compared to *P. spumarius* for all the tested source plants, except alfalfa. It should be noted that acquisition rate of *P. spumarius* from these source plants was lower than those previously registered from olive (≈20-50%); similarly, inoculation success was higher onto periwinkle following acquisition from periwinkle and milkwort (≈45%) compared to those olive-to-olive (30-35%) (Cornara et al. 2017, 2019, Cavalieri et al. 2019). Sharpshooter vectors in the New World, although tested on different bacterium-plant combinations, showed usually high transmission rates from both experimental plants and artificial diets. *Graphocephala atropunctata* (Signoret) and *Homalodisca vitripennis* (Germar), following acquisition from source plants, showed 90 and 20-30% transmission efficiency, respectively. *Bucephalogonia xantophis* (Berg) showed a transmission efficiency of 15% when fed on source plants and 44% when fed on *Xfm* artificial diet (Hill and Purcell 1995, Daugherty and Almeida 2009, Esteves et al. 2018).

The results of the present studies highlight that *C. viridis* is unlikely to become a relevant vector of *Xf* in Europe. The observed acquisition and especially the inoculation rates are lower than those registered for other competent vectors, both spittlebugs and sharpshooters. Also, it should be noted that *C. viridis* presents some ecological constraints that could further hinder its vector competence, specifically: i) it is mainly present in continental Europe and only seldom in Mediterranean areas, where it is limited to humid areas, and ii) feeding host plants are monocots (mainly *Juncus* spp., *Carex* spp. and Cyperaceae), not known hosts for any *Xf* isolate (Kunz et al. 2010, EFSA Panel on Plant Health (PLH) 2015, Cornara et al. 2019, Delbianco et al. 2021). However, in this study we tested a single *Xf* strain and few host plant species. Transmission capability should be explored also for other pathosystems, made of different *Xf* genotypes and different host-plants, these latter more likely to occur in the agroecosystems where this sharpshooter is more abundant (e.g. vineyard of temperate regions) (Pavan and Gambon 2004, Hasbroucq et al. 2020, Markheiser et al. 2020, Bodino et al. 2021). Also, *C. viridis* oviposits on some woody plants (e.g. *Prunus* spp.), and thus, some feeding events could occur on these plants, representing a potential threat to stone fruits for the transmission of *Xf* subsp. multiplex (Cornara et al. 2019, Greco et al. 2021).

In summary, *C. viridis* is a competent vector of *Xfp* ST53 in laboratory studies, although its role in spreading the bacterium is expected to be negligible, given the low transmission efficiency observed in the present study and its ecological constraints (e.g. ecological niche and host-plant range). Further studies should be carried out to explore its potential in other *Xf* pathosystems within vineyard and stone fruit orchard agroecosystems located in humid areas.

## Acknowledgments

The authors wish to thank Sara Primiceri (CNR-IPSP Torino) for the technical support in the experiments, and Francesco Palmisano and Antonella Saponari (Premultiplication Center, CRSFA Basile Caramia) for the production of the recipient plants. The authors acknowledge Vivaio Regionale «Li Foggi» – Agenzia Regionale attività Irrigue e Forestali (ARIF) and Enzo Manni (SOC. AGR. COOP. ACLI – Racale) for the use of the rearing and transmission facilities.

This project has received funding from the European Union’s Horizon 2020 research and innovation programme under grant agreement No. 727987 “Xylella fastidiosa Active Containment Through a multidisciplinary-Oriented Research Strategy XF-ACTORS”.

## Authors’ Contributions

conceptualisation and methodology, N.B., V.C., M.S., C.D., D.B.; statistical analysis, N.B; investigation, N.B., V.C., C.D., G.A.; resources, M.S., D.B.; writing–original draft preparation, N.B.; writing–review and editing, N.B., V.C., M.S., C.D., D.B.; supervision, D.B.; project administration, M.S., D.B.; funding acquisition, M.S., D.B.

