## Supplementary figures and images for "Transmission of *Xylella fastidiosa* subsp. pauca ST53 by the sharpshooter *Cicadella viridis* from different source plants and artificial diets"

### Supp. Figure S1

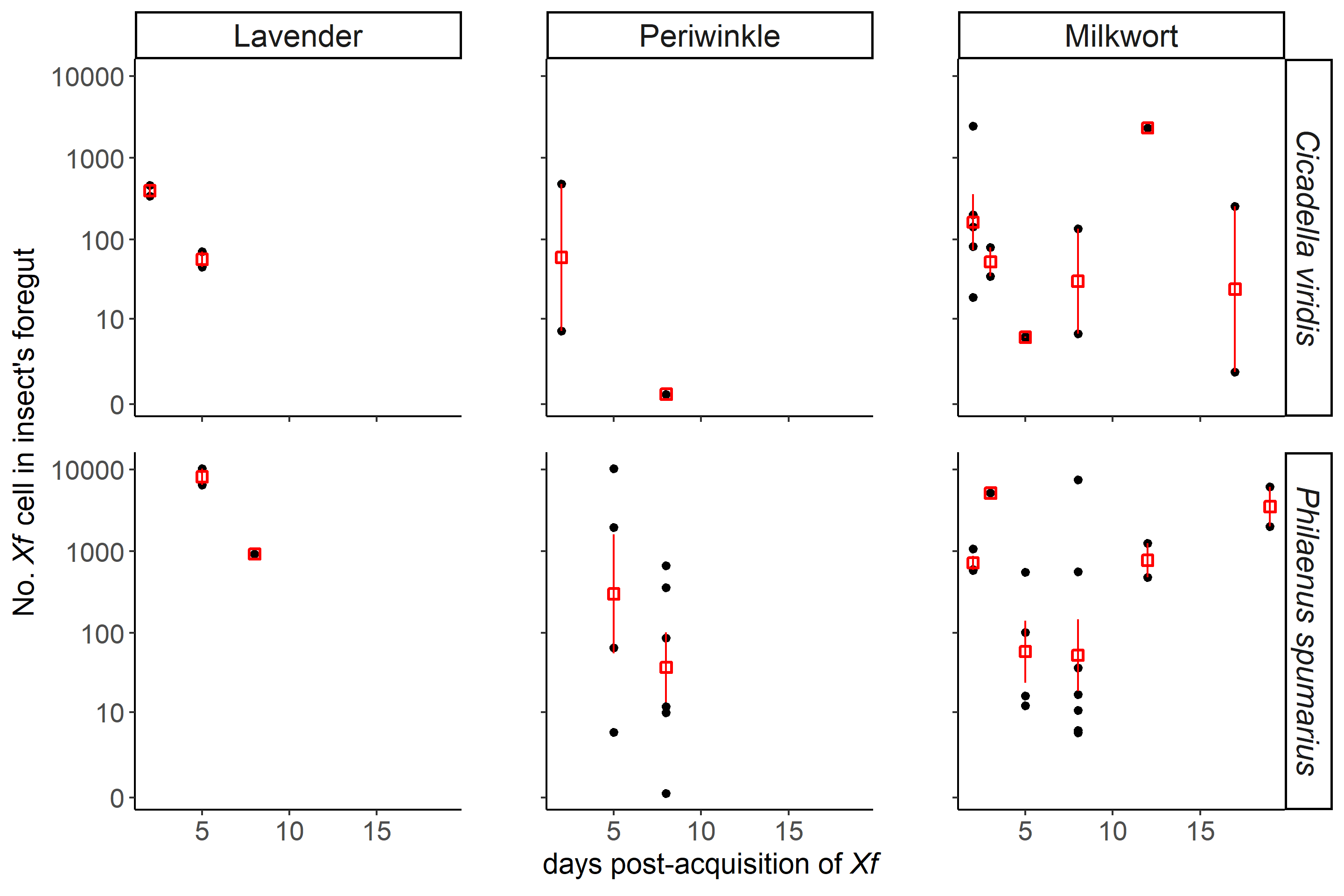
